# LimeSeg: A coarsed-grained lipid membrane simulation for 3D image segmentation

**DOI:** 10.1101/267534

**Authors:** Sarah Machado, Vincent Mercier, Nicolas Chiaruttini

## Abstract

Bioimage analysis is an important preliminary step required for data representation and quantitative studies. To carry out these tasks, we developed LimeSeg, an easy-to-use, efficient and modular 3D image segmentation method. Based on the idea of SURFace ELements, LimeSeg resembles a highly coarse-grained simulation of a lipid membrane in which a set of particles, analogous to lipid molecules, are attracted to local image maxima. The particles are self-generating and self-destructing thus providing the ability for the membrane to evolve towards the contour of the object of interest. We characterize the emergent mechanical properties of this system and show how it can be used to segment many 3D objects from numerous types of image of biological samples (brain MRI, cell epithelium, cellular organelles). LimeSeg is available as a Fiji plugin that includes simple commands, a 3D visualizer, and customization options via ImageJ scripting.

## Background

Over the recent years tremendous improvements have been made on the techniques to acquire 3D images of biological samples at various scales. The volumetric datasets that can be acquired by optical and electron microscopy, as well as with magnetic resonance imaging (MRI), broaden the scientific questions that can be investigated. As a part of this process, it became a standard practice to extract and analyze objects morphology in 3D. However delimiting complex objects such as cells and organelles within a 3D image stack is often a very challenging and time-consuming task. Many tools already exist [1] that can be broadly classified in the following categories: intensity-based methods (region-growing), mathematical morphology (watershed) [2, 3], active contours [4, 5, 6, 7], level-set [8], gradient vector flow [9], machine learning (pixel classification, deep learning)[10, 11].

We developed a new 3D segmentation method, called LimeSeg (Lipid Membrane Segmentation) which has been optimized to achieve a very good balance between efficiency, accuracy and ease of use. LimeSeg is a revisited version of the active contour method that revives early image segmentation methods based on the coupling of surfels, which are oriented particles reprensenting elements of surface [12].

LimeSeg is applicable to segment 3D object coming from an extensive range of sources (magnetic resonance imaging, focused ion beam scanning electron microscopy (FIB-SEM), light microscopy). Additionally, multiple surfel systems can be coupled to segment simultaneously non-overlapping objects, resembling watershed method in that aspect. LimeSeg is implemented as an ImageJ / Fiji [13, 14, 15] plugin, a software which is under very active development and with which the microscopy and image analysis community are already familiar with. It has been optimized to enable segmentation of relatively large images, using graphical processing units for the most consuming time steps and the generic ImgLib2 library [16]. On the user interface side, it can be used with simple pre-defined commands that require initial seeds and 2 parameters. LimeSeg can also be customized by using its recordable graphical user interface (GUI) and the scripting capabilities of Fiji.

## Implementation

### Mathematical description

LimeSeg can be seen as a strongly coarse-grained modeling of a lipid membrane. In this method, a surface is defined by a set of oriented particles (also previously called surfel for SURFace ELement), analogous to lipid particles. The set of surfels evolves until collocating with the surface of a 3D object, labeled on its outline. The number of surfels is adapted locally to allow for area changes subsequent to surface deformation. Surfel displacement is ruled by surfel interactions with the underlying 3D image and by short-range pair interactions between surfels.

Each surfel *i* is defined by a position in 3D: **p_i_** and by a unit normal vector: **n_i_** (Fig. 1A). The rules controlling the interactions between neighboring surfels were chosen based on the segmentation stability, consistency, reproducibility and speed, unlike more physically meaningful simulations [17, 18, 19]. In agreement with previous works for oriented particle systems [12], we found that considering only pairs interaction, short-range coupling with a few layers of neighboring surfels, and three interactions that are detailed below (**F_dist_**, **F_plane_**, **T_tilt_**) were sufficient to fulfill our requirements. Each surfel is also attracted by the local underlying 3D image maximum (**F_signal_**) and is biased by a constant pressure (**F_pressure_**), allowing for surface adaptation to 3D objects contained in the image.

**Figure 1.**
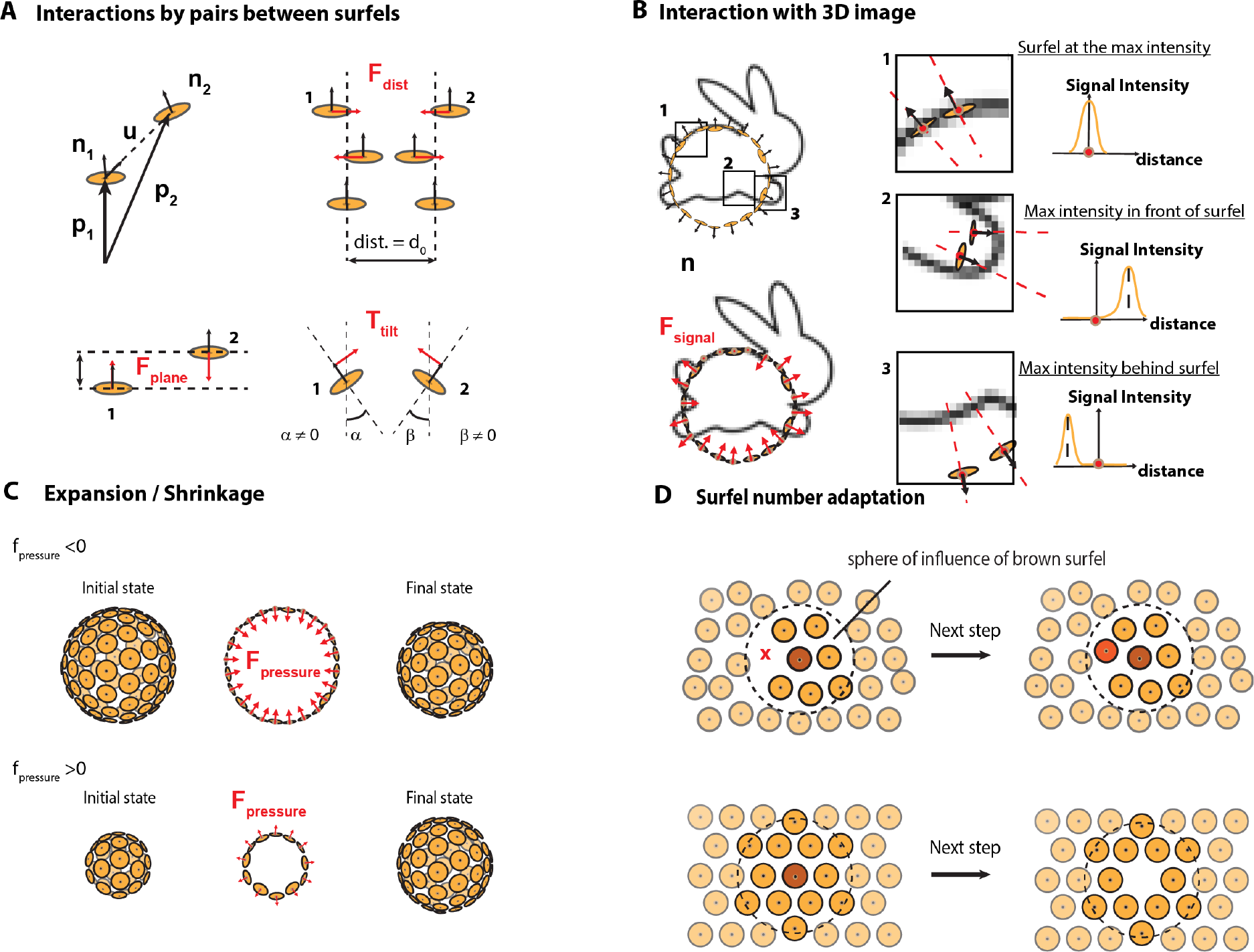
Surfel interaction rules. A ‒ Forces and torque acting on a neighboring pair of surfel. Top left: notation convention for the position and normal vector of surfels. Top right: preferred distance interaction *d*_0_ and associated force **F_dist_**. Bottom left, planar interaction **F_plane_**. Bottom right, **T_tiit_**. B - Interaction with the 3D image. **F_signal_** has a constant norm. It is positive, null, or negative depending on the local image maximum. C - **f_pressure_** exerted along the normal vector. The sign of *f*_*pressure*_ controls surface shrinkage or expansion. D - Adaptation of surfel number depending on local neighbors. The number of neighbors within the sphere of influence is counted. Depending on this number, the surfel is removed or a new one is generated.

Before starting the segmentation, the user has to set the equilibrium distance between pairs of surfels, expressed in units of pixels: *d*_0_. It is the most essential parameter of LimeSeg as it sets the minimal feature size that can be segmented (typically 1 to 10 pixels). The segmentation process then starts from one or several seeds, each seed being a surfel system that delimits a surface. As detailed in the discussion, a seed can be a sphere, a complex shape made from a skeleton, or a pre-existing surfel system. The segmentation is an iterative process that ends when all the surfels have converged, each iteration being divided into 6 phases that are detailed below:

#### 1 Neighboring surfel identification

Each surfel has a fixed-radius sphere of influence. At this step, each surfel identifies and counts the number of surfels comprised within its sphere of influence. We name these surfels neighbors. The radius of the sphere of influence is by default 1:75×*d*_0_.

#### 2 Neighboring surfel forces computation (Fig. 1A)

Surfels interact with neighbors comprised through pair interactions. We note *d* = ‖**p_j_** − **p_i_**‖ the distance and **u** = (**p_j_** − **p_i_**)/*d* the unit vector between surfels *i* and *j*. The first pair interaction **F_dist_**_*j*→*i*_ = *f* (*d*/*d*_0_)**u** maintains the preferred distance *d*_0_ between pairs of surfels. If two surfels are separated by a distance smaller than *d*_0_, **F_dist_** is a harmonic repulsive force. If the distance between the pair is above *d*_0_, **F_dist_** mimics a bond that can break: it is attractive, vanishes at *d* = *d*_0_ and with *d* → ∞. The second interaction **F**_**plane**__*j*→*i*_ = *k_plane_*(**u** · (**n_i_** + **n_j_**))**n_i_** and third pair interactions **T**_**tilt**__*j*→*i*_ = *k_tilt_*(**n**_i_ · **u**)**u** act on the position and the surfels normal respectively. They both favor particle co-planarity.

#### 3 Interaction with the image

To deform the segmented surface so that it adapts to the underlying 3D image, two additional forces are exerted along the normal direction of each surfel. First, **F_signal_** = ±*f_signal_***n**, is a data attachment term of constant norm that links the particles to the 3D image. The direction of this force depends on the local image maximum location relatively to the normal vector (Fig. 1B). Second, **F_pressure_** = ±*f_pressure_***n**, is a fixed global pressure set by the user. This pressure tends to induce the shrinking or the expansion of the surface (Fig. 1C). It is equivalent to the “balloon force” used in related segmentation method [6, 20]. If the surfel is located near to a local image maximum, both **F_signal_** and **F_pressure_** become null.

#### 4 Surfel number adaptation

During segmentation, the number of surfel needs to adapt: the number of surfel has to diminish while the surface shrinks and increase during surface expansion. For local surfel number adaptation, we implemented the following rules (Fig. 1D): 1) If the number of surfels comprised in its sphere of influence is higher than an upper limit, the surfel removes itself. 2) If the number is smaller or equal to the lower limit, a new surfel is created at the position of lowest surfel density. Practically, this position is estimated by computing the sum of the repulsive forces exerted by neighboring surfels. We implemented an equilibration period of a few iterations during which a newly created surfel can neither disappear nor generate a new neighboring surfel. Finally, to allow for clearance of spurious surface, surfels that are too isolated to create new surfels (because their number of neighbors is below the threshold) are removed.

#### 5 Update of surfel position and orientation

The numerical integration follows an explicit Eulerian scheme combined to a purely viscous behavior: at each integration step, the displacement of each surfel is equal to its resulting force multiplied by *d*_0_: **p_i_**(*t*+1) = **p_i_**(*t*)+*d*_0_×[Σ_*j∈neighbors*_(**F**_**dist**__*j*→*i*_+**F**_**plane**__*j*→*i*_)+**F_pressure_**+**F_signal_**], and the normal of each surfel is summed with the resulting sum of torques **n_i_**(*t* + 1) = **n_i_**(*t*) + Σ_*j∈neighbors*_**T**_**tilt**__*j*→*i*_. With this integration scheme, forces can be directly interpreted as displacement per integration step, in units of *d*_0_. For instance, if a constant force of value 0.01 is exerted on a surfel, and if *d*_0_ is set to 15 pixels, it will require 100 steps to move the surfel by 15 pixels.

#### 6 Convergence test

The iterative process stops when all surfels are locked, as they met two convergence criterions. First, each surfel is considered as having converged when it undergoes little displacement or rotation in the course of a defined number of integration steps. Second, when all its neighbors have converged, the position and normal vector of a defined surfel are locked. The above two-step convergence can be used to restrict the active computation zone and speed up the segmentation (see FIB-SEM segmentation part).

### Implementation efficiency

Of the different phases of the integration, neighboring surfels identification, (also called fixed-radius near neighbor search problem), is the most computationally intensive. To accelerate this step, we implemented a custom space-partitioning tree building algorithm, which is further parallelized on graphics processing units (GPU) using CUDA library and the Java JCUDA wrapper. The computation of pairs of forces is also a time consuming step which can also be processed on GPU. Nonetheless CPU computation remains faster for low number of particles, thus LimeSeg automatically switches between CPU and GPU with a threshold at 20k surfels. Overall, with these optimizations, an integration step scales almost linearly with the number of particles (*N*^1.05^), and three parts (1-2-3) contributes almost equally to 90 % of the integration time.

## Results

We first review the emergent properties of the simulated set of particles. Understanding the surface behavior is critical when it comes to segmenting 3D images.

### Leakage / Arrest

A common problem to overcome is leakage: the surfel surface improperly spread through small “holes” where the data outline is missing or weak. It artificially includes into the object voxels that not do belong to the object. Conversely, little holes could also be part of an object (the beginning of a “neck” or of a tube for instance). In that case, the surfel system has to go through the hole. Indeed, a surface that would stop around the hole would converge without reaching the outline of the objet, resulting in an incomplete segmentation. To understand how LimeSeg behaves and ultimately be able to tune LimeSeg correctly, we illustrate how LimeSeg segments a test case consisting of a 100×100×100 image cut in half by a plane containing a circular hole of radius *r*_*hole*_ in its center. We segment this image while varying the radius of the hole *r*_*hole*_ and *f*_*pressure*_ and keeping *d*_0_ constant. Depending on these parameters, two outputs are observed: 1 - the surface stops around the hole (required in the case of artefactual holes in the image signal) or 2 - the surface goes through the hole and continues growing (required to segment an object containing a tube, a neck) (Fig. 2A). In a diagram plotting *r*_*hole*_/*d*_0_ as a function of *f*_*pressure*_, these two segmentation outcomes are in two distinct domains separated by a 1/r curve. In other terms, the frontier is governed by an intrinsic quantity which is equal to the radius multiplied by the pressure. This quantity has the unit of a surface tension, and reveals an intrinsic emerging threshold of the system. It can be understood as follows: during segmentation, the radius of the hole combined with the applied pressure sets transiently a surface tension that can be computed by the Young-Laplace equation and that is withheld by the particle network. If the tension is above the threshold, the links between surfels are disrupted and new surfels are generated to fill the gaps, leading to expansion of the surface. If the tension is below the threshold, surfel interactions are maintained, the surface is stable and the convergence can be reached without going through the hole. In conclusion, the user can tune *f*_*pressure*_ and do to adapt the segmentation to the required output. Intuitively, lowering *d*_0_ or increasing *f*_*pressure*_ leads to a better penetration of the surface through gaps.

**Figure 2.**
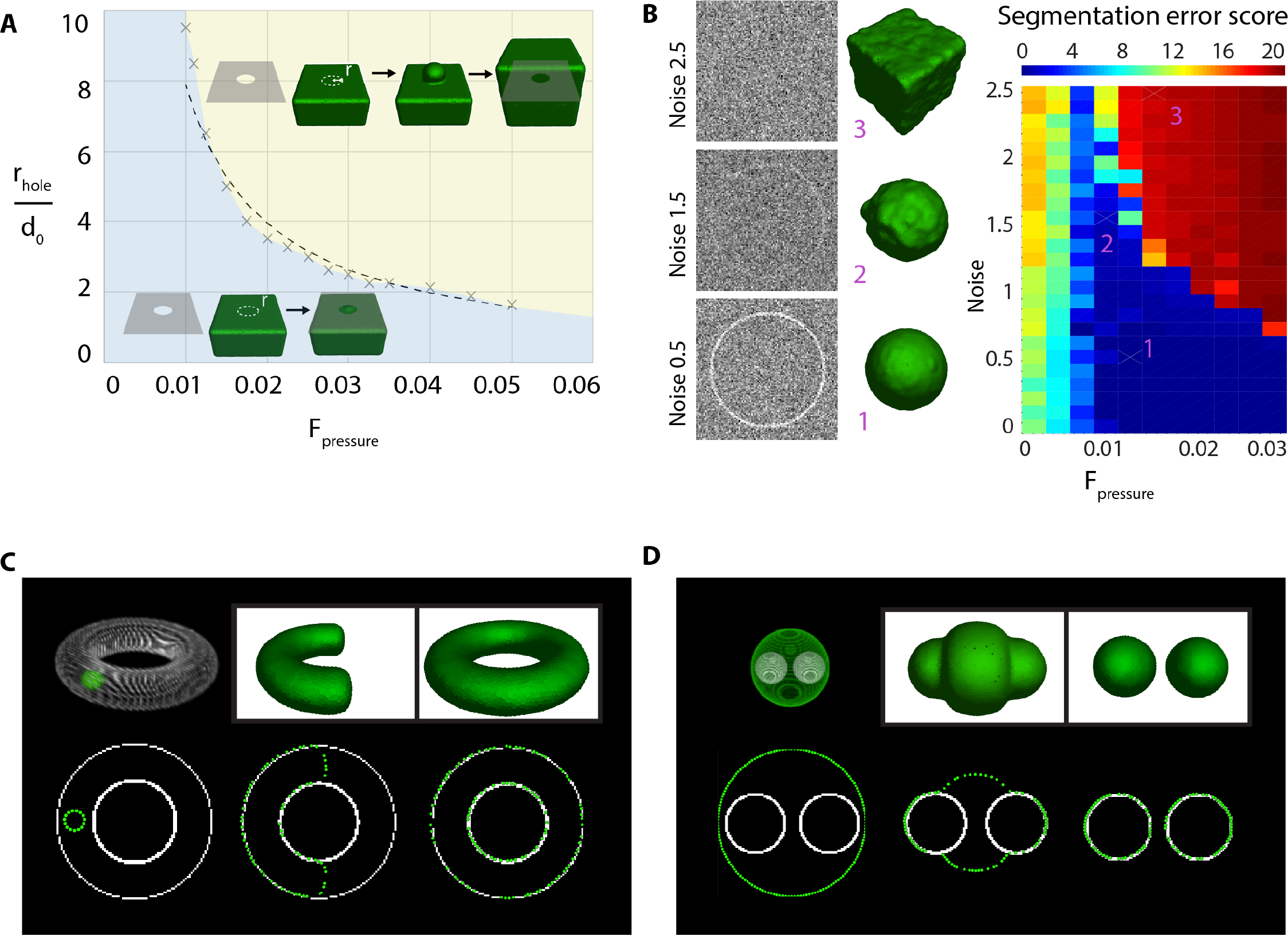
Point cloud mechanics characterization. A - Behavior of surfels network with fixed do as a function of *f*_*pressure*_ when it encounters a circular hole of radius *r*_*hole*_. In the blue region, the surfel mesh does not cross the hole. In the yellow region the surfel mesh flows through the hole. A 1/r dotted line approximates the frontier between these regions. B - Image noise segmentation benchmark, see text for details. Left: equatorial plane of sphere image with various noises and resulting segmentation. Right: Segmentation score (i.e. root mean square of surfel distance to the target sphere in pixel) as a function as noise and *f*_*pressure*_, all other parameters are unchanged. C - Surface fusion test. The initial state consists of a spherical seed inside a torus. After several iterations, the shape of the segmentation surface successfully merges with itself (*f*_*pressure*_ > 0). D - Surface fission test. The initial state consists of a spherical seed surrounding two spherical objects. The segmentation surface successfully splits during the course of the segmentation (*f*_*pressure*_ < 0).

### Noise resilience

Another common matter of interest in segmentation is the method resilience to image noise. We address in a simple test case how the signal to noise ratio affects the segmentation outcome. Starting from a slightly off-centered spherical seed, we segment a larger sphere while varying the noise contained in the image. In our test image, the edge of the sphere has on average 2 pixels in thickness, with a variability depending on the 3D rasterization, and a signal value of 1. A Gaussian noise centered on zero was added on the image, with standard deviation values ranging from 0 to 2.5 (typical lateral slices are shown Fig. 2B, left). We evaluated the segmentation outcome with positive *f*_*pressure*_ values varying from 0.005 to 0.035. To evaluate the reliability of the segmentation, we measured after convergence, the deviation of the distance from each particle to the target sphere (Fig. 2B, right). For positive pressure of too low intensity (< 0.01), even a very small noise was preventing the seed inflation. It leads to incorrect segmentation. Conversely, the seed could pass the outline of the sphere on a signal to noise ratio dependent manner in the case of high positive pressures (> 0.03). There is an optimal value for *f*_*pressure*_ around 0.01, where the outline can be detected at a low signal to noise ratio. For noisy images, a *f*_*pressure*_ value around 0.01 is thus recommended.

### Surface topology

Another important information to know for a segmentation method is how it handles topological changes, i.e. can a surface spontaneously split and merge? To test LimeSeg for intrinsic merging, we segmented a torus starting from a spherical seed located inside the torus. We used a positive pressure and observe that the two ends of the “C” shape are fusing to form the torus (Fig. 2C). To test for fission, we segmented two spheres starting from one unique spherical seed, which was surrounding the two target spheres. We applied a negative pressure and starting from one seed we obtained two distinct segmented surfaces (Fig. 2D). In conclusion, LimeSeg handles topological changes such as fusion and fission. In contrast with mesh methods, the topological changes naturally arise from the particle set interaction rules.

## Discussion

LimeSeg user interface is composed of a recordable GUI, an ImageJ1 application programming interface, and provides a 3D viewer. LimeSeg is intended to be used to segment one or multiple surfaces, in a modular fashion. Unlike most segmentation tools, it generally does not require any image pre-processing.

Below, we demonstrate the capabilities and versatility of LimeSeg. We segment lipid vesicles, a human brain, the plasma membrane and endoplasmic reticulum of a HeLa cell and the cells of a Drosophila egg chamber. The imaging modalities (confocal microscopy, MRI, FIB SEM), image contrast and resolution, signal to noise ratio, shape size and objet density are also different in these examples. With these test cases we cover a broad range of image segmentation challenges in field ranging from developmental biology to biomedical imaging.

### Segmentation of lipid vesicles

The first example of application consists of segmenting the surface of deformed lipid vesicles which are attached on a glass coverslip. The vesicles are imaged with a confocal microscope that outputs a 3D image stack. We segmented two vesicles sequentially, starting from spherical seeds located inside each vesicle. We show in Fig. 3 how the point set matches the outlines of these two vesicles. Each vesicle segmentation takes a few seconds, and each vesicle is made of approximately 1000 particles.

**Figure 3.**
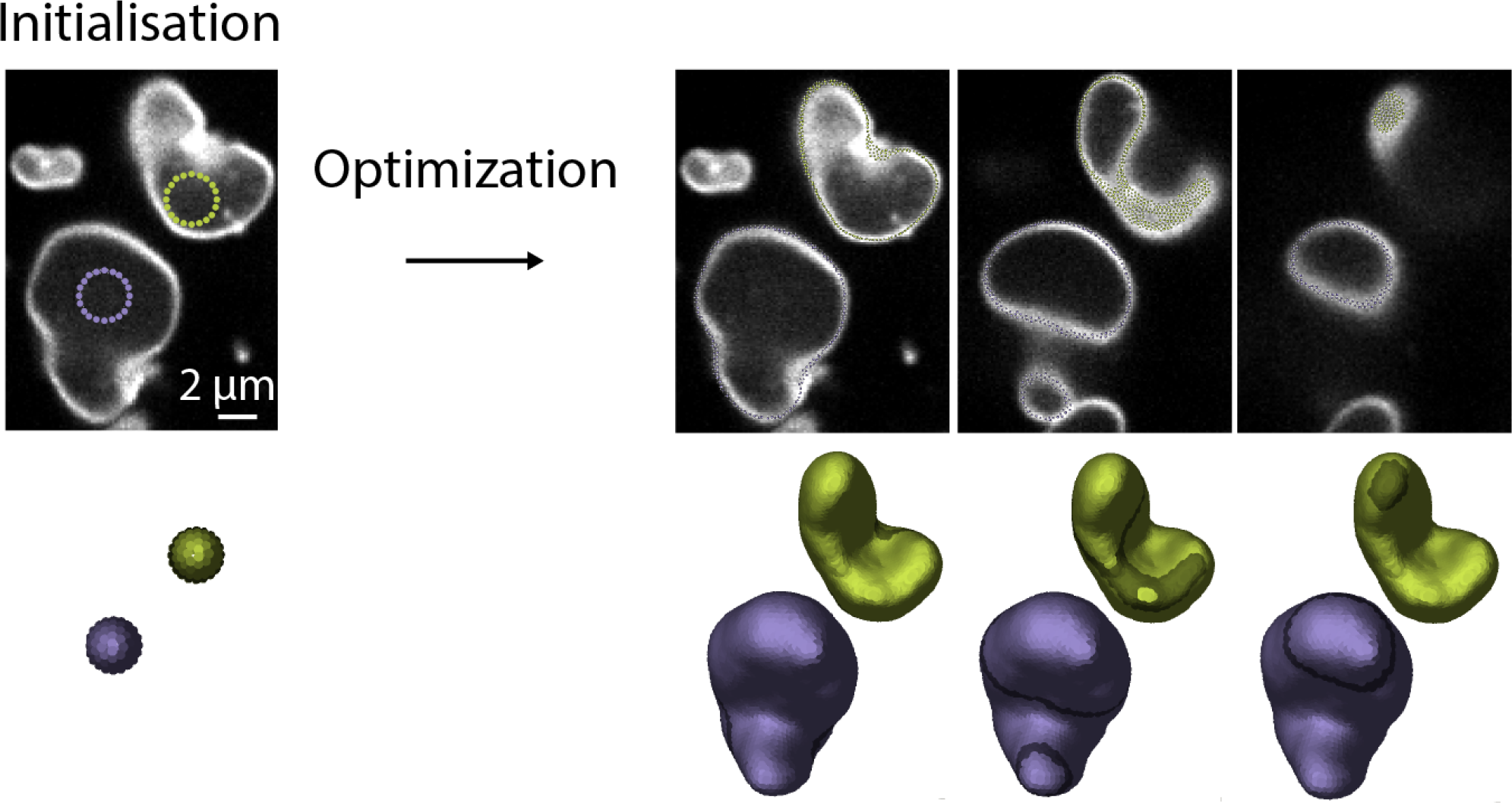
Segmentation of deformed lipid vesicles. The two vesicles are segmented sequentially. Right: segmentation outcome. Three z slices where surfels appear as dots are shown as well as the 3D reconstruction, where the in-planes surfels are highlighted.

### Segmentation from MRI images: full human brain segmentation

In this test case, we segment the cortical surface of an MRI dataset (FLAIR sequence, see Fig. 4, bottom). The dataset consists of 512×512×224 voxels, and we set an initial “skeleton” seed which is slightly larger than the brain. This skeleton (or non-spherical seed) is used to initialize the segmentation and consists of roughly defined ROIs surrounding the brain at specific slices through the stack (Fig. 4, left). Using the user specfied *d*_0_ value, the plugin can dispatch surfels on this basic geometrical skeleton before starting the segmentation. When convergence is reached with *d*_0_ = 4, one can notice that finer details of the cortex are missed: like in the blue region of Fig. 2A, the radius of circumvolutions are too small relatively to *f_pressure_* and *d*_0_. In such a case, the segmentation process can be refined by progressively varying *d*_0_, starting from the previous final conditions. We refined the segmentation by reducing *d*_0_ down to 1.5 pixels and resumed the segmentation to reach final convergence. At the end of the segmentation, the fine brain circumvolutions are detected, (Fig. 4, right). The whole process took 5 minutes and resulted in a 300 000 points cloud.

**Figure 4.**
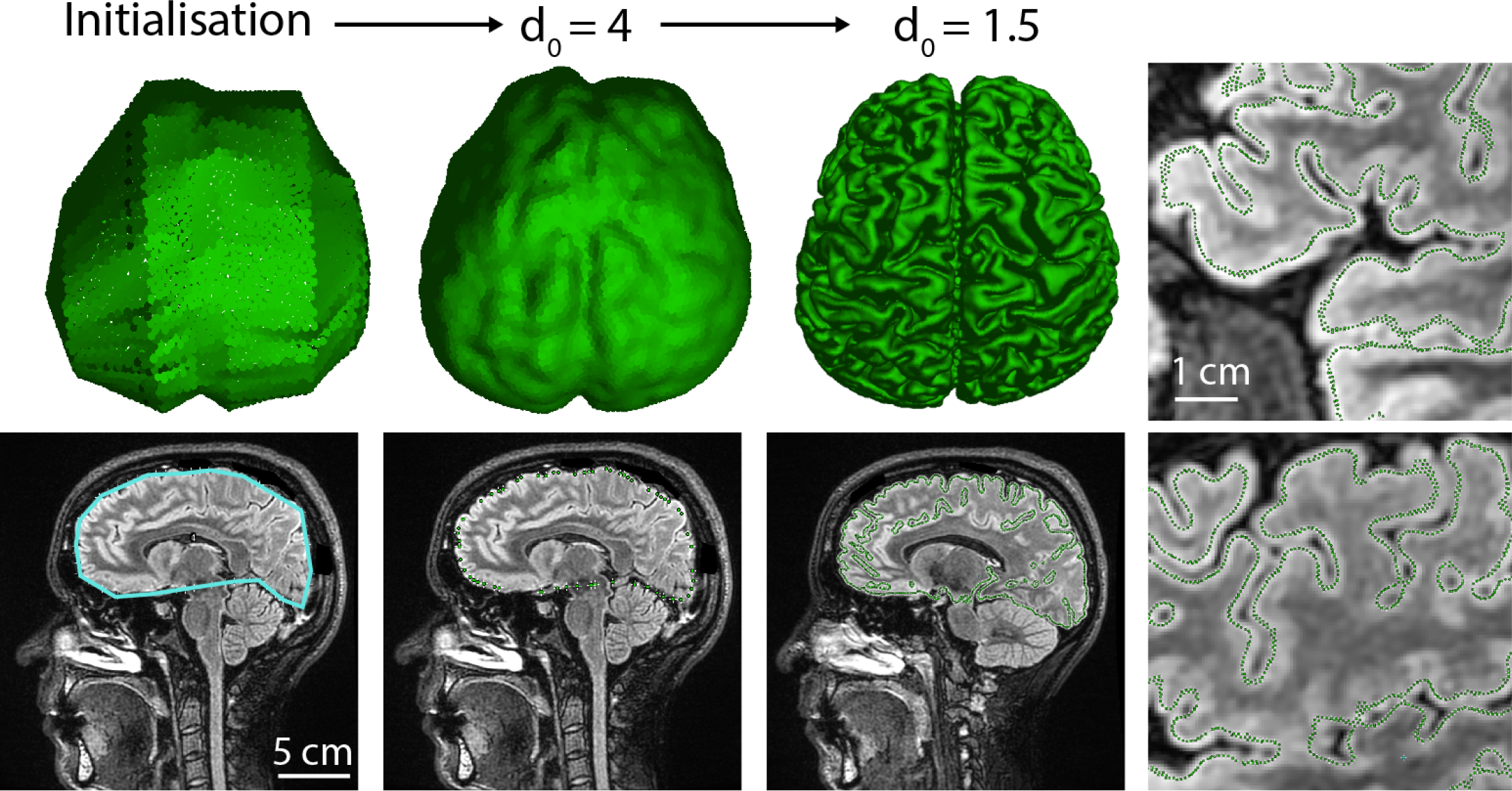
Human brain MRI surface segmentation. From left to right: 1 - initialization of the shape with ROI skeleton (blue line on the data image). 2 - After segmentation convergence with *d*_0_ = 4, many details of the cortex are missed. 3 - Segmentation refinement by decreasing *d*_0_ to 1.5. 4 - Zooms showing details being retrieved by the finest segmentation where surfels appear as green dots.

### Plasma membrane and endoplasmic reticulum segmentation

We challenged the method by segmenting a 3D EM dataset of 4300×4100×650 voxels. The dataset consists of nearly isotropic sections of a Hela cell (4.13 nm in XY, 5 nm in Z) (Fig. 5A) prealigned with TrackEm2 [21], without additional preprocessing. We aimed to segment two structures sequentially: the plasma membrane and the endoplasmic reticulum (ER). For such a large dataset, it is expected that during the course of the segmentation, a large portion of surfels will have converged while relatively small actively growing regions will continue to grow. Keeping all the points that have converged at each integration step induces unnecessary computational costs. To circumvent this problem, LimeSeg, like other segmentation methods [22], has a way to restrict the computation to actively segmenting regions. In brief, while constructing the space partitioning tree, LimeSeg detects and replace large chunks of locked surfels by a single super surfel. The conversion of active surfels into passive and locked super surfels is reversible. If a particle that is not locked is interacting with a super surfel, the super surfel is replaced by the chunk of surfels it contains in the next integration step, allowing for rearrangements.

The plasma membrane segmentation was performed starting with 5 spherical seeds located outside of the cell. The seeds inflated and merged until surrounding the cell. This segmentation took 4 hours and resulted in a 4 million particle point cloud (Fig. 5B, 5C, green). We next segmented the ER system. We initiated the segmentation with 5 spherical seeds located into the lumen of the ER and ran 80000 integration steps over 6 hours. This led to a cloud of 15 million points in which the double nuclear envelope, that is inherently linked to the ER system, was segmented as well (Fig. 5B, 5C, magenta). Some limitations can be seen: inexistent holes are sometimes detected and the ER cannot be segmented when the double membrane is too thin (Fig. 6D). We believe that this segmentation is still very satisfactory given the very little amount of work required by the user. Many aspects of the cell membranes geometry are preserved and can be detected in the segmentation: membrane invaginations like clathrin coated pits (Fig. 5E), nuclear pore complexes (Fig. 5F), the complex network of intertwined filopodia and the highly circumvoluted ER shape (Fig. 5C). Thus, the segmentation generated with LimeSeg provides a very good starting point for further shape analysis, like proper surface quantification and curvature measurements.

**Figure 5.**
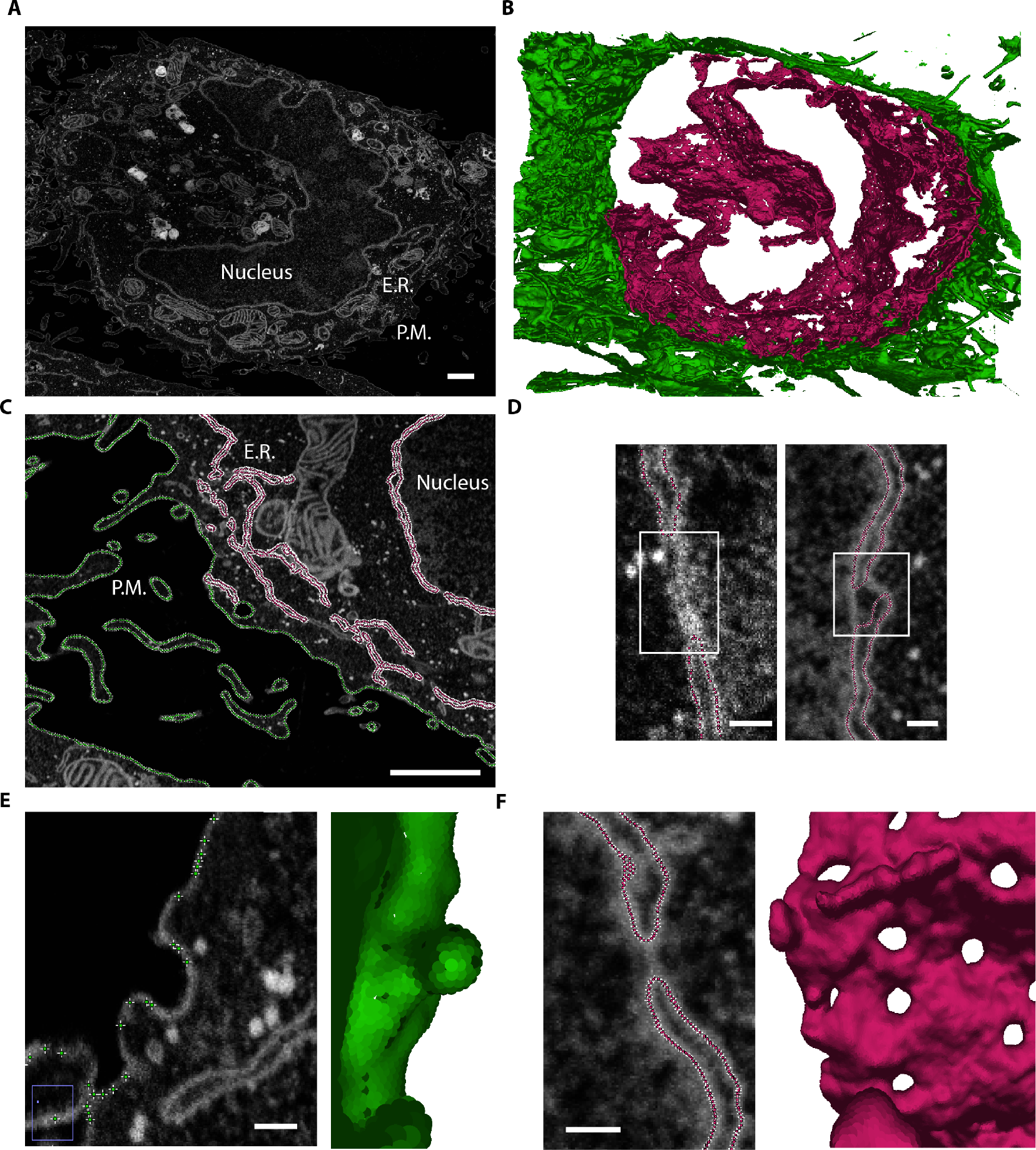
Endoplasmic reticulum (ER) and plasma membrane (PM) segmentation of a FIB-SEM dataset. A - Typical data slice where the nucleus, E.R. and P.M. are visible. B ‒ Resulting segmentation of E.R. (magenta) and the plasma membrane (green). C - Segmented ER and PM, overlaid on the original data. D - Missed parts of nuclear envelope where it is too thin to be correctly segmented (left), spurious hole generated during segmentation (right). E - Detail showing Plasma membrane invagination in 2D and 3D. F - Detail of nuclear pore complex as seen on 2D and on 3D. Scalebars: A, C: 1 *μm*; D, E, F: 100*nm*

**Figure 6.**
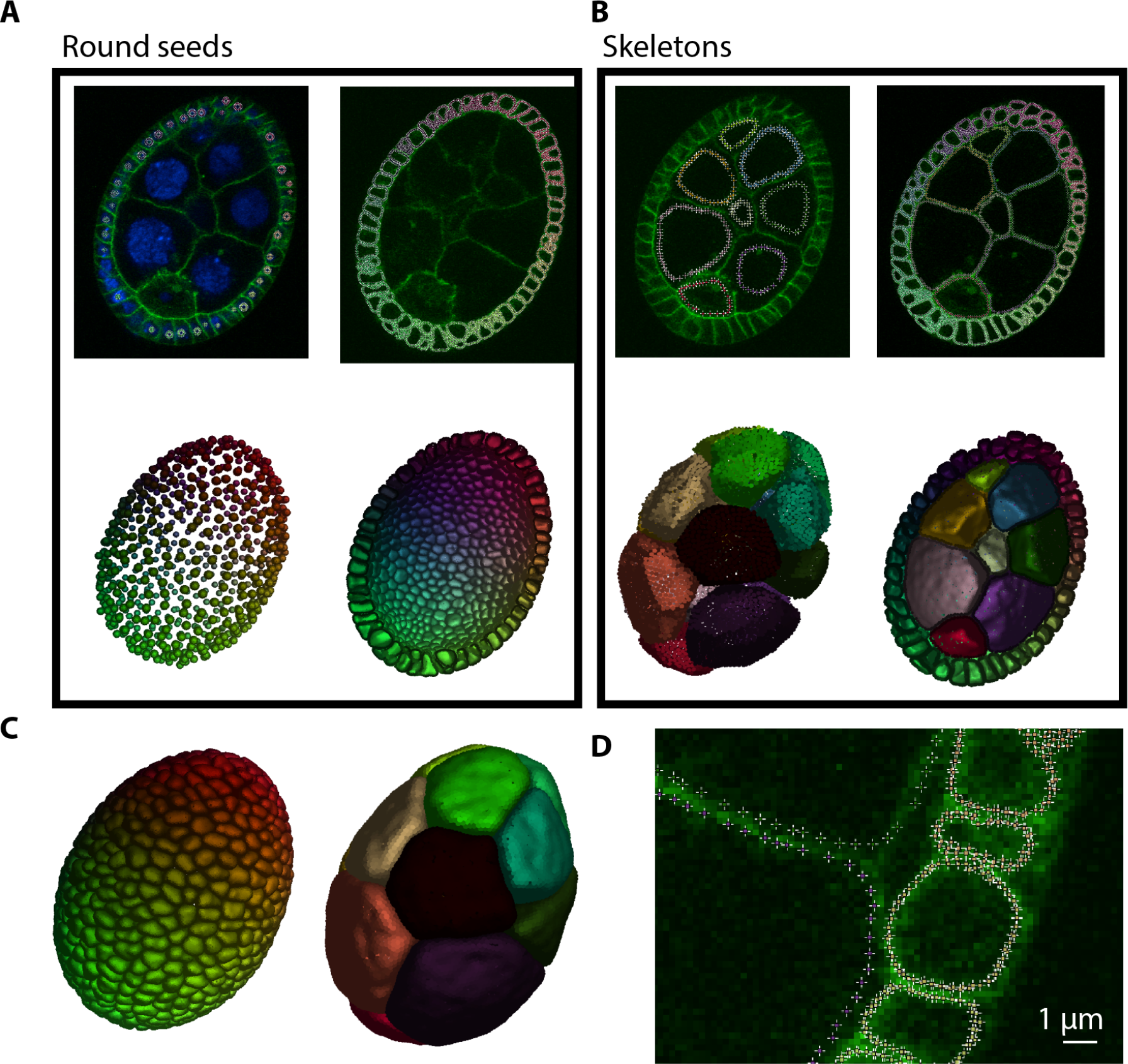
Drosophila ovary segmentation. A - Segmentation of the follicle cells. Up: surfels appear as colored dot (one color per cell). Below: cut of 3D reconstruction, where only the surfels below the showed slice are shown. Left is the initial state and right shows the state after convergence. B - Segmentation of the nurse cells. During the course of segmentation, surfels of follicle cells are added and locked to maintain the outline. C - 3D reconstruction output for nurse cells and follicle cells. D - Detail of surfel position found after segmentation.

### Cell segmentation and cell volume measurement form confocal images: the case of a Drosophila egg chamber

In the previous examples, only one object is being segmented at the time. We show with this example that multiple objects can be segmented simultaneously by delimiting cells from confocal fluorescent slices of a drosophila egg chamber (Fig. 6A). The egg chamber is an interesting case study as it consists of three different types of cells which shape and size are very different: the nurse cells, the follicles cells and the oocyte. As a prerequisite for the segmentation, the user needs to provide LimeSeg the approximate position of each cell. This seeding can be done in many ways, it can be manual or by identifying local minima in a blurred image, or, like we did, by using the fluorescent channel of nuclei and by computing the barycenter of each nuclear blob. At each position, a sphere of predefined radius serves as an initial point set. The user then specify that each sphere is a different object and then LimeSeg sets a unique identifier to all surfels of a particular cell spherical seed. This part is in contrast with the the FIB-SEM dataset, where each sphere was attributed to a unique object identifier. Based on these identifiers, surfel-surfel interactions are differentiated: if two interacting surfels belong to the same cell, the interactions are as described before but if two surfels of different cells interact, they are not considered as neighbors, and only the repulsive part of **F_dist_** is kept, allowing for surfaces repulsion. We first segmented the external layer of the ovary using a low value for *d*_0_, necessary to resolve the geometry of these small cells ( 5 minutes) (Fig. 6A). Then we locked this set of points, which define the external layer of the embryo, and we initialized the 16 big cells with a higher *d*_0_ value and skeleton seeds (Fig. 6B, left). These 16 cells are then segmented while keeping fixed points of the small cells to maintain the outline (Fig. 6B, right). We show in Fig. 7C the segmentation result for these two types of cells, and in Fig. 7D some details of surfel positioning in the image. The processing of this example image took 10 minutes and resulted in a cloud of 380 000 points, which can be used for further quantifications.

### Cell tracking and shape analysis of LimeSeg outputs

Even if not shown in this manuscript, LimeSeg supports analysis of time series and multichannel images. In particular, for limited shape changes of objects between successive frames, the object shape can be segmented over time by providing the output of the previous frame as an input to the following frame.

A point cloud is the structure used during segmentation, but it is not the best structure objects to perform further object shape analysis. A polygonal mesh is much more suited. LimeSeg provides a surface reconstruction algorithm from its point cloud, and functions for basic shape analysis (volume / surface / surface center of mass). The data can be accessed directly via ImageJ commands, or via scripting. As an alternative to ImageJ, the point cloud and/or the meshes can also be exported in the standard ply file format, which then can be imported into other software for further processing and analysis.

More detailed instructions regarding the software usage, customization, tutorials improvements and updates are available on the ImageJ wiki (https://imagej.net/LimeSeg).

## Future directions and conclusion

Potential future development includes a better integration with ImageJ2 and with the expected new generation of FIJI 3D viewer that is still under development. On the algorithmic side, several improvements can be envisioned for instance by modifying the interaction of dots with the underlying images. Particularly appealing is the idea of modifying surfels interaction with a filtered gradient vector image.

In conlusion, we implemented a new surface reconstruction plugin adapted for various sources of images, which is deployed in the user-friendly and well-known ImageJ environment. It requires little to no image pre-processing. The plugin is available via its ImageJ update site (http://sites.imagej.net/LimeSeg/) and the code is available on GitHub (https://github.com/NicoKiaru/LimeSeg) under MIT license.

## Author’s contributions

NC devised the segmentation method. SM, NC and VM generated experimental and theoretical datasets. NC and SM implemented the software and designed and conducted the analyses. SM and NC wrote the manuscript.

## Acknowledgements

The authors thank Emmanuel Derivery, Serge Dmitrieff, Marcos González-Gaitán, Jean Gruenberg, Rita Mateus, Aurélien Roux, Daniel Sage and Caterina Tomba for support throughout this project and critical comments of the manuscript, the Fiji community for the help regarding ImageJ2 integration. The authors thank the PFMU facility of the University of Geneva (https://www.unige.ch/medecine/pfmu/en/) for help with SEM/FIB.

